# Dispersal syndromes affect ecosystem functioning in ciliate microcosms

**DOI:** 10.1101/2021.02.19.431939

**Authors:** Allan Raffard, Julie Campana, Delphine Legrand, Nicolas Schtickzelle, Staffan Jacob

## Abstract

Dispersal is a key process mediating ecological and evolutionary dynamics. Its effects on metapopulations dynamics, population genetics or species range distribution can depend on phenotypic differences between dispersing and non-dispersing individuals (*i.e*., dispersal syndromes). However, scaling up to the importance of dispersal syndromes for meta-ecosystems have rarely been considered, despite intraspecific phenotypic variability is now recognised as an important factor mediating ecosystem functioning. In this study, we characterised the intraspecific variability of dispersal syndromes in twenty isolated genotypes of the ciliate *Tetrahymena thermophila* to test their consequences for biomass productivity in communities composed of five *Tetrahymena* species. To do so, dispersers and residents of each genotype were introduced, each separately, in ciliate communities composed of four other competing species of the genus *Tetrahymena* to investigate the effects of dispersal syndromes. We found that introducing dispersers led to a lower biomass compared to introducing residents. This effect was highly consistent across the twenty *T. thermophila* genotypes despite their marked differences of dispersal syndromes. Finally, we found a strong genotypic effect on biomass production, confirming that intraspecific variability in general affected ecosystem functions in our system. Our study shows that intraspecific variability and the existence of dispersal syndromes can impact the functioning of spatially structured ecosystems in a consistent and therefore predictable way.

## Introduction

Dispersal is a complex and key process mediating ecological and evolutionary dynamics in spatially structured landscapes (Mouquet and Loreau 2003, Bowler and Benton 2005, Ronce 2007, Edelaar et al. 2008, Clobert et al. 2012, Edelaar and Bolnick 2012). Spatially structured ecosystems (*i.e.*, meta-ecosystems) strongly rely on dispersal rates (*i.e*., the proportion of dispersing individuals, Loreau et al. 2003b) as dispersal mediates species interactions and nutrient fluxes in ecosystems, which in turn modifies ecosystem functions (Loreau et al. 2003b, Harvey et al. 2016). In meta-ecosystem modelling, dispersing individuals are mostly considered as phenotypically similar to non-dispersing ones, although the dispersal status of individuals (*i.e.*, being a disperser or a resident) is often associated with strong phenotypic specializations (Bowler and Benton 2005, Clobert et al. 2009). Especially, the willingness and ability of organisms to disperse at each of the three dispersal phases (*i.e.*, emigration, transience, and immigration) is often associated with various phenotypic traits including behaviours, morphology, life history or physiology (Clobert et al. 2009). Such suite of traits differing between dispersers and residents, named dispersal syndromes, have been described in many taxa (Clobert et al. 2009, Le Galliard et al. 2012, Stevens et al. 2014, Legrand et al. 2015). For instance, dispersers are often larger than residents in freshwater fish (Comte and Olden 2018).

While often differing among species (Stevens et al. 2014, MacLean and Beissinger 2017), dispersal syndromes can also vary within species depending on both evolutionary and ecological factors (Bowler and Benton 2005, Legrand et al. 2016, Cote et al. 2017a). For instance, genetic covariances between dispersal and reproductive traits are found in wing-dimorphic crickets (reviews in Zera and Brisson 2012, Saastamoinen et al. 2018). In toads, interaction between landscape matrix composition and predation risk shapes movers phenotype (Winandy et al. 2019). Therefore, variability in genetic composition and/or local environmental conditions (biotic or abiotic) can generate variability in dispersal syndromes among populations (Cote et al. 2017a). In turn, not only dispersal rate itself, but also dispersal syndromes (regardless of whether they differ within species or not) can have consequences on ecological dynamics (Jacob et al. 2019). Some studies showed that phenotype-dependent dispersal can be important in mediating metapopulation dynamics, species interaction, and phenotypic range distribution (Duckworth and Badyaev 2007, Phillips et al. 2010, Shine et al. 2011, Messager and Olden 2019). In ciliate microcosms, dispersal syndromes involving morphological and demographic traits can affect metapopulation size and stability (Jacob et al. 2019). The consequences of dispersal syndromes at the scale of ecosystems have however rarely been studied, despite the fact that phenotypic variability within species can affect multiple ecosystem functions, including nutrient recycling, decomposition rate and primary production (Harmon et al. 2009, Bassar et al. 2010). For instance, variability in sticklebacks (*Gasterosteus aculeatus*) morphological traits (*e.g.*, mouth apparatus and body shape) can affect primary production and carbon cycle through difference in diet among individuals (Harmon et al. 2009, Schmid et al. 2019). Importantly, such intraspecific variability can affect ecosystem functions as much as species richness (Des Roches et al. 2018, Raffard et al. 2019).

The impacts of dispersal syndromes and their intraspecific variability on ecosystem functioning could be due to variability of interaction strengths in communities between dispersers and residents. First, dispersers and residents could differ in their competitive abilities, in turn modifying species coexistence (Hart et al. 2016, Turcotte and Levine 2016). Some traits often involved in dispersal syndromes, such as aggressiveness, may shape competition strength in colonised habitat and affect species coexistence (Duckworth and Badyaev 2007, Wolf and Weissing 2012), which could affect ecosystems. Dispersal syndromes can also modulate ecosystem functioning through trophic interactions when resource use differs between residents and dispersers (Cote et al. 2017b). Hence, primary evidence emerged, showing that dispersers of *Dikerogammarus villosus* -a freshwater amphipod decomposer- consume more detritus than residents, increasing the decomposition rate of organic material in colonizing habitats (Little et al. 2019). Interestingly, effects differed between the two related study species, suggesting that the effects of dispersal syndromes on ecosystem functioning might be species-dependent. Since current rapid environmental changes are shaping variability within species and notably in dispersal syndromes (*e.g.*, habitat fragmentation; Cote et al. 2017a), unexpected effects going along with alteration of the dispersal process could modify ecosystem functioning.

Here, we aimed at testing whether dispersal syndromes and their intraspecific variability affect ecosystem functioning. We used microcosms to quantify the effects of dispersal syndromes of multiple isogenic strains (hereafter referred to as *genotypes*) of the ciliated protist *Tetrahymena thermophila* on biomass production of communities composed of four competing species of the genus *Tetrahymena*. As a prerequisite to this study, we quantified dispersal syndromes of *T. thermophila* and their variability among 20 genotypes: for each genotype, we measured morphological traits (*i.e.*, cell size and shape) and demographic parameters (*i.e.*, population growth and maximal density) of resident and disperser individuals. We then tested whether residents and dispersers differently affect biomass production, while controlling for density. We further tested whether intraspecific variability in dispersal syndromes (*i.e.*, differences between genotypes in the measured dispersal-related traits) mediates these ecosystem effects. We expected dispersal syndromes to affect biomass production because of functional differences between dispersers and residents (Little et al. 2019), and we expected that these effects might be genotype-dependent if the traits associated with dispersal vary among genotypes.

## Material and methods

### Study species

Native from North America, *Tetrahymena thermophila* is a 30–50μm unicellular eukaryote living in freshwater ponds and streams (Collins 2012, Doerder and Brunk 2012). Twenty isogenic genotypes (strains D1 to D20, characterised in Pennekamp et al. 2014, Table S1) were used to assess variability in dispersal syndromes. All genotypes displayed clonal reproduction in our culture conditions (Elliott and Hayes 1953, Bruns and Brussard 1974). Cells were maintained in axenic rich liquid growth medium (0.5% Difcoproteose peptone, 0.06% yeast extract) at 23°C (Schtickzelle et al. 2009, Altermatt et al. 2015) in sterile conditions under a laminar flow hood.

### Quantification of dispersal syndromes

Dispersal syndromes, and their dependency to genotypes, have already been described in microcosms of *T. thermophila* (*e.g.*, Fjerdingstad et al. 2007, Jacob et al. 2019a, 2019b). We built on these previous studies to quantify them at a wider intraspecific level and test for their effect at the ecosystem level. Dispersal rates and syndromes were quantified using standard two-patch systems (*e.g.*, Schtickzelle et al. 2009, Pennekamp et al. 2014, Jacob et al. 2019b). They consist in two habitats (1.5 ml microtubes) connected by a corridor (4 mm internal diameter, 2 cm long silicone tube) filled with growth medium. We measured dispersal rate and characterised resident and disperser cells in five replicated systems for each genotype. To do so, cells from each genotype were placed in one patch at a standardised density (40,000 cells/ml) during 30 minutes of acclimation, and the corridor was then opened to allow cells to disperse for four hours towards the initially empty neighbour patch. Corridors were then closed to prevent further movements until measurement. Population growth rate is low enough during such a four hour time frame in this species to avoid bias in the quantification of dispersal rates (Pennekamp et al. 2014, Jacob et al. 2018).

Five samples of 10μl were pipetted in each of the two patches (for resident and disperser cells) and pictured under dark-field microscopy (Axio Zoom V16, Zeiss) to assess cell density and morphology using a procedure developed within the ImageJ^®^ software (Pennekamp and Schtickzelle 2013). First, the dispersal rate for each genotype was calculated as 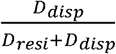, where *D_disp_*, *D_resi_* are dispersers’ and residents’ abundances respectively. Second, *cell size* (cell surface area on pictures) and *cell elongation* (aspect ratio, *i.e.*, ratio of cell major/minor axes) were estimated in both residents and dispersers.

Then, demographic traits (*i.e.*, growth rate and maximal population density) of residents and dispersers of each genotype were measured. To do so, ~150 cells of residents or dispersers (mean volume ± SE: 14.02 ± 1.25 μl) from each patch of each dispersal system were transferred into 96-well plates (250μl wells) filled with growth medium, with two technical replicates (averaged in subsequent analyses). Absorbance at 450nm was recorded every two hours for two weeks using a microplate reader (Infinite 200, TECAN) to quantify population growth. We used a general additive model (R-package *gam*; R Core Team 2013, Hastie 2018) to smooth the observed absorbance measurements to avoid any bias due to slight variability in absorbance measures, and fitted logistic growth curves using the *grofit* package (gcfit function with spline fit; Kahm et al. 2010). *Growth rate* was then assessed as the maximum slope of population growth, and *maximal population density* as the density reached at the plateau (Jacob et al. 2017).

### Experimental communities and ecosystem functioning

Competitive communities were assembled using four competing *Tetrahymena* species to which residents or dispersers of each *T. thermophila* genotype from the first part of experiment were added. A single strain of each competing species was used (*T. americanis* A5, *T. borealis* B8, *T. pyriformis* P4, and *T. elliotti* E5; Table S1) to avoid potential differences of competitive interactions due to genetic variability. To assemble communities, we transferred ~150 cells of each of the four competing species (10 μl of cultures previously diluted to 15,000 cells/ml) into 96-well plates (250 μl wells filled with growth medium) and then introduced dispersers/residents from one specific *T. thermophila* genotype (~150 cells, see above). In total, we assembled 200 communities with equal initial abundance of all species: 20 *T. thermophila* genotypes × 2 dispersal status (resident and disperser) × 5 replicates. For each replicate of the dispersal experiment (see above), we additionally run two technical replicates of community dynamics that were averaged for analyses. Absorbance at 450nm was recorded every two hours for two weeks to quantify the dynamics of ecosystem functioning by measuring the biomass of the communities in the microcosms. As for individual genotypes growth (see above), we fitted logistic growth curves to absorbance data and computed *biomass production* as the maximum slope of community absorbance increase, and the *maximal biomass* as the maximal absorbance reached by the community.

### Statistical analyses

We primarily aimed at confirming that morphological traits (*i.e.*, cell size and cell elongation) and demographic parameters (*i.e.*, growth rate and maximal density) differed between residents and dispersers in *T. thermophila* and that these differences vary among genotypes, as observed in our previous studies. Firstly, population growth rate, maximal density, cell size and elongation were summarised in a Principal Component Analysis (PCA, *ade4* R-package, Chessel et al. 2007). Secondly, we fitted two linear models (lm function, *stats* R-package) with the scores on the first two PCA axes (either PCA1 or PCA2) as dependent variables, and the dispersal status (*i.e.*, disperser or resident), genotype identity and their interaction as explanatory variables.

To test for the impact of dispersal syndromes and genotype identity on ecosystem functioning, we fitted linear models with biomass production or maximal biomass as dependent variables, and dispersal status (residents *vs* dispersers, two levels factor), genotype identity, and their interaction as explanatory variables. Then, we fitted similar models by replacing genotype identity and the dispersal status by the scores of PCA1 and PCA2 axes summarising phenotypic traits, in order to test whether the dispersal syndrome (*i.e.*, the measured phenotypic variability underlying differences between dispersers and residents) and genotype identity affect biomass.

## Results

The first two axes of the PCA on resident and disperser phenotypic traits explained 58% and 26% of phenotypic variation, respectively. The first axis (hereafter named *demographic axis*) was positively associated with population growth rate (loading = 0.90) and maximal density (0.92), with high values on this axis referring to cells with high growth capacities (Figure 1). The second PCA axis (hereafter named *morphological axis*; Table S2) explained variance in cell size (0.64) and elongation (−0.57), with high values describing larger and rounder cells. The twenty genotypes of *T. thermophila* strongly differed in their position on both the demographic and morphological axes (Table 1). Additionally, genotypes differed in the intensity and direction of the resident-disperser differences, both in terms of demography and morphology (significant genotype × dispersal status interactions; Table 1). This means that the described dispersal syndrome varied among genotypes in this species. We also detected a small difference of morphology between residents and dispersers independently of genotype identity, residents being slightly smaller overall (Table 1).

**Figure 1.**
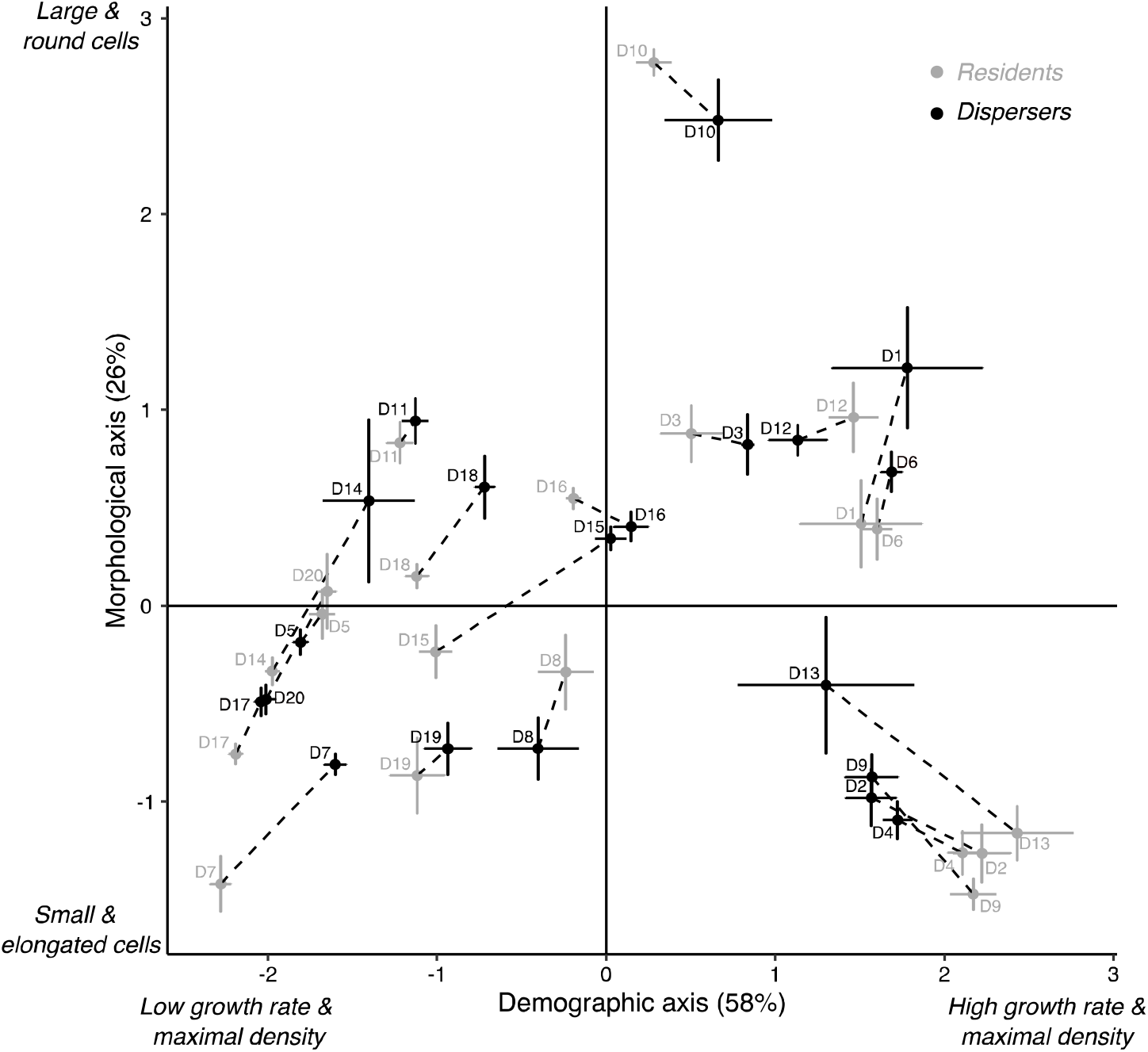
Variability of phenotypic traits (extracted from PCA) among twenty genotypes of *Tetrahymena thermophila*. The first axis expresses demography (population growth rate and maximal density), and the second axis expresses morphology (cell size and elongation). Mean values (±SE among 5 replicates) for residents and dispersers for each genotype are displayed.

**Table 1.** Effect of dispersal status (dispersers vs residents) and *T. thermophila* genotype on the position in the PCA space defined by a demographic axis and a morphological axis.

|  | Sum sq | df | F | *p*-value | R<sup>2</sup> |
| --- | --- | --- | --- | --- | --- |
| Demographic axis |  |  |  |  | 0.94 |
| Dispersal status | 0.07 | 1, 160 | 0.448 | 0.504 | 0.00 |
| Genotype | 431.16 | 19, 160 | 138.025 | <0.001 | 0.91 |
| Dispersal status × Genotype | 12.50 | 19, 160 | 4.003 | <0.001 | 0.03 |
| Morphological axis |  |  |  |  | 0.90 |
| Dispersal status | 2.23 | 1, 160 | 17.903 | <0.001 | 0.01 |
| Genotype | 178.05 | 19, 160 | 75.180 | <0.001 | 0.85 |
| Dispersal status × Genotype | 8.09 | 19, 160 | 3.419 | <0.001 | 0.04 |

*T. thermophila* dispersers and residents differed in their impacts on biomass (Figure 2). Specifically, introducing dispersers of any *T. thermophila* genotype led to communities with lower biomass production and maximal biomass than introducing residents of the same genotype at the same density (Figure 2). This effect of dispersal status on biomass production and maximal biomass was highly consistent across genotypes (non-significant dispersal status × genotype interactions; Table 2a). The genotype × dispersal status interaction still has a marginal effect on biomass production (Table 2a), suggesting that some differences among genotypes might mediate the effects of residents and dispersers on biomass production. Additionally, independently of being a resident or a disperser, cells from different *T. thermophila* genotypes affected biomass production and maximal biomass in communities, suggesting an effect of intraspecific variability on ecosystem functioning (Figure 2 and Table 2a).

**Figure 2.**
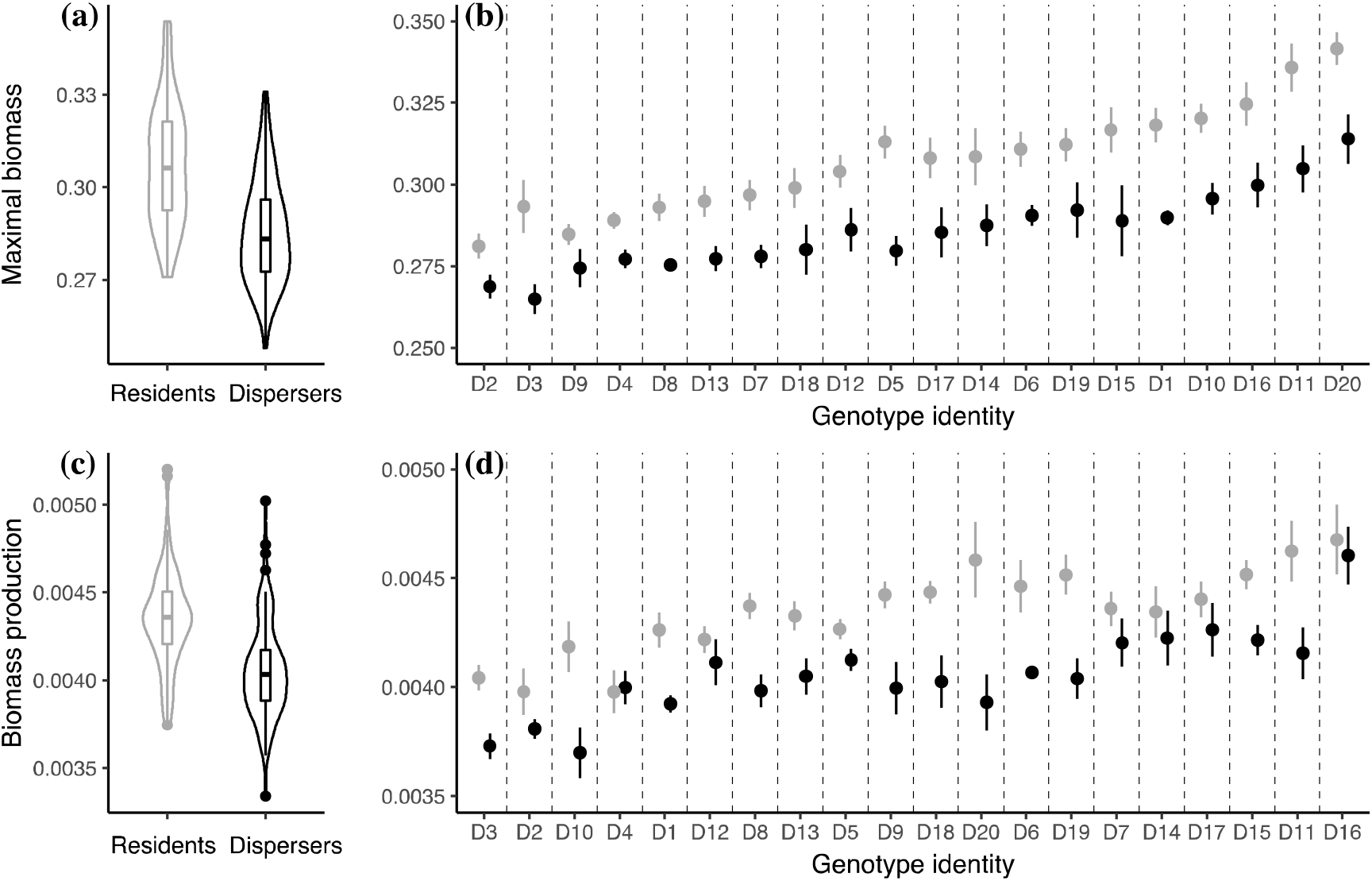
Impact of dispersers and residents on community biomass: effects of introducing *T. thermophila* residents or dispersers of each genotype in a ciliate community on its maximal biomass **(a & b)** and biomass production **(c & d)**. Left panels **(a & c)** are averaged effects over all 20 genotypes, whose individual results are presented on right panels **(b & d)**, sorted from small to high mean.

**Table 2.** Effect on maximal biomass and biomass production of the 40 types of *T. thermophila* cells that were introduced (20 genotypes × 2 residents/dispersers status) considered either as combinations of the dispersal status and genotype identity **(a)** or as positions that they occupy along the demographic and morphological PCA axes **(b).** Significant variables are in bold.

|  | Sum sq | df | F | *p*-value | R<sup>2</sup> |
| --- | --- | --- | --- | --- | --- |
| **(a)** |  |  |  |  |  |
| Maximal biomass |  |  |  |  | 0.69 |
| Dispersal status | **0.023** | **1, 160** | **140.766** | **< 0.001** | **0.26** |
| Genotype | **0.036** | **19, 160** | **11.567** | **< 0.001** | **0.41** |
| Dispersal status × Genotype | 0.002 | 19, 160 | 0.615 | 0.891 | 0.02 |
| Biomass production |  |  |  |  | 0.60 |
| Dispersal status | **4.2x10<sup>-6</sup>** | **1, 160** | **88.839** | **< 0.001** | **0.21** |
| Genotype | **6.2x10<sup>-6</sup>** | **19, 160** | **6.831** | **< 0.001** | **0.31** |
| Dispersal status × Genotype | 1.4x10<sup>-6</sup> | 19, 160 | 1.593 | 0.063 | 0.07 |
| **(b)** |  |  |  |  |  |
| Maximal biomass |  |  |  |  | 0.15 |
| Demographic axis | **0.007** | **1, 197** | **19.898** | **< 0.001** | **0.09** |
| Morphological axis | **0.005** | **1, 197** | **29.909** | **< 0.001** | **0.06** |
| Biomass production |  |  |  |  | 0.07 |
| Demographic axis | **1.3x10<sup>-6</sup>** | **1, 197** | **13.661** | **< 0.001** | **0.06** |
| Morphological axis | 1.4x10<sup>-8</sup> | 1, 197 | 1.521 | 0.219 | < 0.01 |

Then, we explored the mechanisms underlying the effects of dispersal status and genotypes on biomass by quantifying the effects of phenotypic traits that we found to be involved in dispersal syndromes summarised by the demographic and morphological PCA axes. Overall, since the demographic and morphological traits were highly related to genotypic identity, and slightly to the dispersal status (Table 1), they depicted mainly a genotype effect on biomass and to a lesser extend a dispersal status effect. As expected, the morphological axis was linked to maximal biomass, depicting a positive effect of cell size and elongation on maximal biomass (Table 2b, Figure S1). More surprisingly, maximal biomass and production were negatively affected by demographic traits of *T. thermophila* (Table 2b, Figure S1). The higher the growth rate and maximal density of *T. thermophila* cells, the lower the biomass in experimental communities.

## Discussion

Dispersal is an important process affecting metacommunity and meta-ecosystem dynamics (Loreau et al. 2003b, Massol et al. 2017, Gounand et al. 2018), but the role of dispersal syndromes in these dynamics is poorly understood. In this study, we tested whether dispersal syndromes and their intraspecific variability affect ecosystem functioning in ciliate microcosms. As previously shown in *T. thermophila*, residents and dispersers differed in both morphological and demographic traits, a dispersal syndrome that varied among genotypes (Fjerdingstad et al. 2007, Jacob et al. 2019, 2020). Then using five ciliate species community (*T. thermophila* with four competing species), we showed that introducing dispersers consistently led to lower biomass production and maximal biomass irrespective of genotype-specific dispersal syndromes.

Our results strikingly showed that the dispersal status of an individual leads to consistent effects on ecosystem functioning across multiple genotypes, despite we measured intraspecific variability in the phenotypic differences between residents and dispersers. Specifically, dispersers lead to lower biomass than residents, suggesting that they may hence be functionally dissimilar from residents, leading to different interspecific interactions (Chesson 2000, Mitri and Foster 2013). Dispersal often incurs costs, for instance associated with the development of specific phenotypic attributes or with energy and metabolic demands required for moving among habitats (Bonte et al. 2012, Cayuela et al. 2019). We could primarily expect that costs of dispersal would make dispersers less competitive for other species in the communities, which would have probably increased biomass in microcosms compared to residents (because of relaxed competition strength). At the opposite, we could expect that a decrease of biomass production would come from higher competitive abilities of dispersers resulting in an inhibition of biomass production (Mitri and Foster 2013). Competition may indeed lead to mutual inhibition among competing species (most of the communities contained four species -including *T. thermophila*- at the end of the experiment, and *T. elliotti* was always excluded, see Appendix S1), resulting in low growth rate of individual species and eventually a low biomass production (Mitri and Foster 2013, Ghoul and Mitri 2016). In microbial communities, competitive interactions might be caused indirectly through resource exploitation or directly through cell damages (*e.g.*, chemical toxin) (Ghoul and Mitri 2016). Hence, here, the dispersers of *T. thermophila* might increase the strength of competitive interaction (direct and/or indirect), which may partly explain the negative impact on total biomass in microcosms (Thompson et al. 2005, Sandau et al. 2019). Further investigations might benefit the understanding of these results by assessing the competitive strength of residents and dispersers against each one of the other species.

Additionally, investigating the mechanisms underlying the effects of *T. thermophila* cells on ecosystem functioning, we found that demographic parameters and morphological traits can impact biomass. These traits have been previously found to affect metapopulations dynamics in *T. thermophila* (Jacob et al. 2019). Here, the higher the demographic parameters (*i.e.*, growth rate and maximal biomass in isogenic conditions) of the introduced *T. thermophila* cells, the lower maximal biomass and biomass production. While this might seem contradictory, we could speculate on a trade-off between growth parameters of *T. thermophila* (*i.e.*, proxies for fitness without competition) and competitive abilities (Bohannan et al. 2002, Kneitel and Chase 2004, Mille-Lindblom et al. 2006). Competition is associated with energetic costs, and for example the traits necessary for sustaining higher growth (*e.g.*, efficiency of resource uptake) might come at the expense of other traits (*e.g.*, toxin resistance) allowing to face other species competing for the same resource (Mille-Lindblom et al. 2006). Interestingly, those effects were not due to density *per se* as communities were made of species with equal densities. We rather measured *per capita* effects on biomass productivity. Including density effects, and determining the independent effects of density, dispersal syndromes and phenotypic traits, might bring further insights into the role of varying dispersal strategies in meta-ecosystems functioning.

Demography and morphology only partly explained differences between resident and disperser individuals (up to 5% when cumulating simple and interactive effects, see Table 2), while they differed mainly among *T. thermophila* genotypes. Therefore, they probably account for a small amount of the variance explained by the dispersal status on biomass metrics. This is confirmed by direct effects of dispersers and residents on maximal biomass and biomass production, independently of their demography and morphology (see Table S3, and Figure S2 for a conceptual diagram of our results). Dispersal is a complex process that depends on many phenotypic traits (Clobert et al. 2012). It is then not surprising that we were not able to capture much of the phenotypic variance explaining the disperser/resident differences. It has been shown that swimming capacities have strong influence on *T. thermophila* dispersal rates (Pennekamp et al. 2019) but other traits (*e.g.*, metabolism, resource uptake, activity) should probably underline the general and consistent resident *vs.* disperser effects on the measured ecosystem functions.

The rate of dispersal is important for the functioning of meta-ecosystems, metacommunities, and other ecological networks (Loreau et al. 2003b, Mouquet and Loreau 2003, Massol et al. 2011, Baguette et al. 2013, Thompson and Gonzalez 2017). Yet, resident and disperser individuals are mostly considered as similar in those frameworks. We showed that this is not the case, at least in a ciliate example as residents and dispersers differently affected biomass production, with most probably *per capita* effects (*i.e.*, independent of density). While dispersal is often seen to affect competition through changes in density (Thompson et al. 2020), our study suggests that phenotypic differences are also at play. Such a result might be important for theoretical predictions regarding meta-ecosystems dynamics. For instance, in range expansion areas, we could speculate that differences in competitive abilities between residents and dispersers might, independently of their growth capacities, alter species coexistence in edge habitats. Subsequent diversity-ecosystem functioning relationships might be altered (Thompson et al. 2020). Yet, these results might be species-dependent, since dispersers can be stronger or weaker competitors than residents depending on the species (Anholt 1990, Hanski et al. 1991, Bowler and Benton 2005, Cadotte et al. 2006, Duckworth and Badyaev 2007). Overall, our results need to be generalised to other species in different ecological networks, which could ultimately benefit theories on meta-ecosystem functioning by inferring finer predictive outputs.

To conclude, we found that dispersal syndromes, the differences between resident and disperser individuals, affect ecosystem functioning with dispersers decreasing biomass productivity compared to residents. This effect was not dependent upon the intraspecific variability in dispersal syndromes we have measured in this experiment. These results suggest that individual strategies linked to dispersal can affect the dynamics of meta-ecosystems, potentially through variability in competitive abilities. Measuring other dispersal-related traits than those related to morphology or growth (*e.g.*, behavioural and physiological) would allow assessing more precisely the mechanisms explaining the described effects of dispersal syndromes on ecosystems. An interesting perspective would be to integrate the ecological differences between dispersers and residents in spatially explicit meta-ecosystem models to test for the effects of dispersal syndromes on several aspects of meta-ecosystem dynamics.

## Acknowledgements

We warmly thank Michèle Huet for maintaining cell cultures essential for this study. AR was supported by a F.R.S-FNRS research project (PDR T.0211.19) awarded to NS in collaboration with SJ. NS is Senior Research Associate of the F.R.S-FNRS. SJ and JC acknowledge financial support from the Agence Nationale de la Recherche for the project CHOOSE (ANR-19-CE02-0016). SJ, JC and DL are part of the TULIP (Laboratory of Excellence Grant ANR-10 LABX-41). This paper is contribution of the Biodiversity Research Center at UCLouvain.

