## Supplementary material for "Dispersal syndromes affect ecosystem functioning in ciliate microcosms": Supp_info

**Table S1.** Codes and Tetrahymena Stock Center (https://tetrahymena.vet.cornell.edu) identity (TSC ID) of the 20 genotypes of *Tetrahymena thermophila*, and the single genotype of the four competing *Tetrahymena* species used in the experiment.

| Code | TSC ID | Species |
| --- | --- | --- |
| D1 | SD01546 | *T. thermophila* |
| D2 | SD01547 | *T. thermophila* |
| D3 | SD01548 | *T. thermophila* |
| D4 | SD01549 | *T. thermophila* |
| D5 | SD01550 | *T. thermophila* |
| D6 | SD01551 | *T. thermophila* |
| D7 | AK III | *T. thermophila* |
| D8 | SD01553 | *T. thermophila* |
| D9 | SD01552 | *T. thermophila* |
| D10 | SD01557 | *T. thermophila* |
| D11 | SD01558 | *T. thermophila* |
| D12 | SD01556 | *T. thermophila* |
| D13 | SD01555 | *T. thermophila* |
| D14 | SD01554 | *T. thermophila* |
| D15 | SD01560 | *T. thermophila* |
| D16 | SD01559 | *T. thermophila* |
| D17 | SD01561 | *T. thermophila* |
| D18 | SD01562 | *T. thermophila* |
| D19 | SD01564 | *T. thermophila* |
| D20 | SD01563 | *T. thermophila* |
| A5 | SD03182 | *T. americanis* |
| B8 | SD03155 | *T. borealis* |
| P4 | SD01671 | *T. pyryformis* |
| E5 | SD01674 | *T. elliotti* |

**Table S2.** Loading of each trait on the first two axes of a PCA analysis performed on four traits measured on the 20 genotypes of *T. thermophila*. Bold values represent variables contributing more than 20% to the axis construction (values into brackets).

|  | Axis 1 | Axis 2 |
| --- | --- | --- |
| Maximal density | **0.92 (36.6)** | -0.41 (12.1) |
| Growth rate | **0.90 (34.8)** | -0.35 (16.5) |
| Cell size | 0.56 (13.4) | **0.64 (39.5)** |
| Cell shape | -0.59 (15.2) | **-0.57 (31.9)** |

**Table S3.** We tested whether dispersal status and genotype identity affect directly ecosystem functioning beyond and independently of the effects of traits (morphology and demography). To do so, we fitted linear models including maximal biomass (or biomass production) as dependent variable, and the demographic and morphological axes (PCA axes, see the main text), dispersal status, genotype identity and the genotype × dispersal status interaction as explanatory variables.

|  | Sum sq | df | F | *p*-value | R^2^ |
| --- | --- | --- | --- | --- | --- |
| Maximal biomass |  |  |  |  | 0.70 |
| **Dispersal status** | **0.022** | **1, 158** | **129.759** | **<0.001** | **0.26** |
| **Genotype** | **0.022** | **19, 158** | **6.815** | **<0.001** | **0.35** |
| Dispersal status × Genotype | 0.002 | 19, 158 | 0.518 | 0.952 | 0.02 |
| Demographic axis | < 0.001 | 1, 158 | 0.186 | 0.667 | 0.03 |
| Morphological axis | < 0.001 | 1, 158 | 0.027 | 0.869 | 0.02 |
| Biomass production |  |  |  |  | 0.61 |
| **Dispersal status** | **4.1x10^-6^** | **1, 158** | **84.905** | **<0.001** | **0.21** |
| **Genotype** | **5.1x10^-6^** | **19, 158** | **5.556** | **<0.001** | **0.29** |
| Dispersal status × Genotype | 1.4x10^-6^ | 19, 158 | 1.609 | 0.059 | 0.07 |
| Demographic axis | 7.5x10^-8^ | 1, 158 | 1.578 | 0.211 | 0.03 |
| Morphological axis | 3 x10^-10^ | 1, 158 | 0.007 | 0.936 | 0.003 |

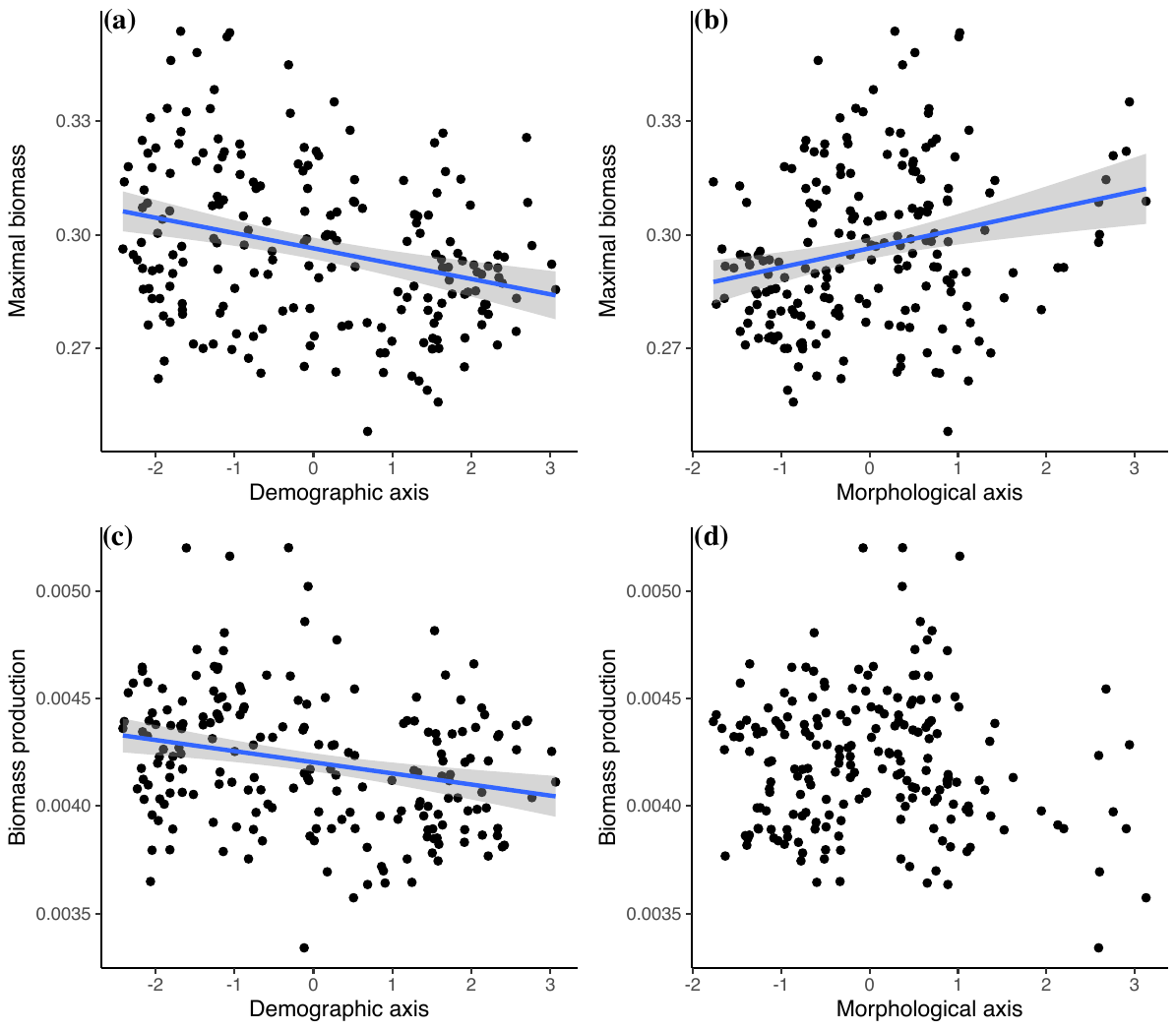

**Figure S1.** Relationships between demographic and morphological axes (PCA axes) with maximal biomass, and biomass production. Lines are displayed for significant relationship (see Table 1 in the main text) and grey shadows represent 95% confident interval.

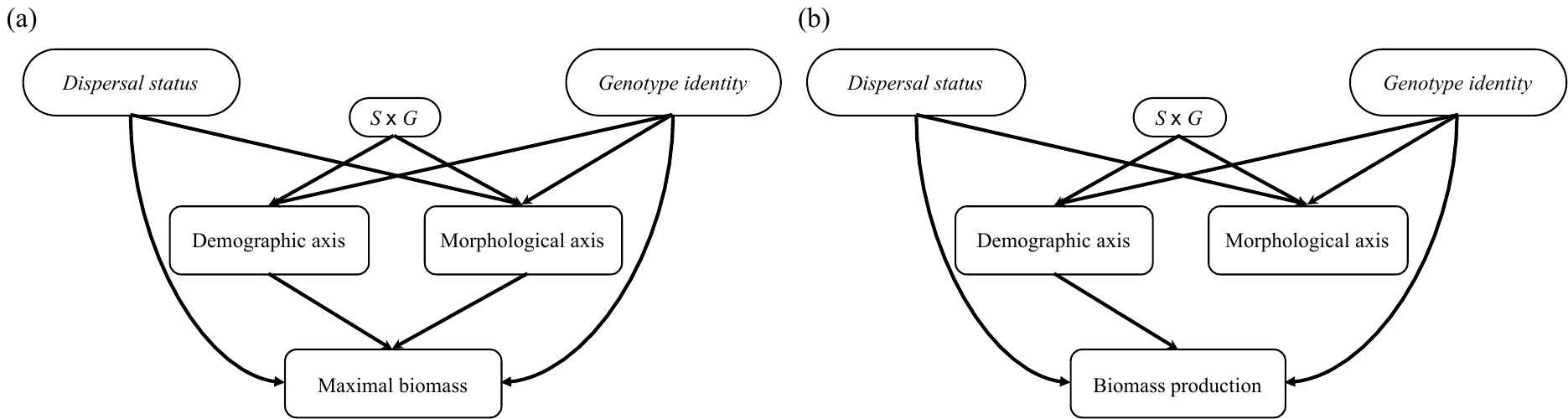

**Figure S2.** Conceptual diagram summarizing our results. It depicted the direct and indirect effects of residents and dispersers (dispersal status *S*) and genotype identity (*G*) on maximal biomass **(a)** and biomass production **(b)**. ‘S × G’ denotes the interaction between the dispersal status and the genotype identity. The indirect effects -although very low compared to direct effects- were supported by the impacts of the demographic and morphological axes (PCA axes, see the main text) on maximal biomass and biomass production.

**Appendix S1**

We performed an additional analysis using flow cytometry (FACS Canto 2, BD Sciences, Yi and Dean 2013, Bestion et al. 2018) to check for the presence or absence of the five species at the end of the experiment. When communities reached the plateau, 10 μl of each sample were analysed with fast flux settings (66 μL min^-1^) to quantify side scatter (SSC), forward scatter (FSC), and four fluorescence parameters: green (FL1), orange (FL2), red (FL3) and blue (FL4). The six metrics were used in a random forest algorithm using the *randomForest* R-package (Liaw and Wiener 2002) to determine species-identity for each cell. Random forest algorithm was previously trained on data obtained from monocultures (*i.e.*, isogenic culture of each competing species; dispersers and residents of *T. thermophila* parallelly grown up in the same conditions), and then applied to assign species identity to each particle detected by the flow cytometer (mean estimated error rate ± SD of the random forest discrimination: 0.018 ± 0.006). We did not detect any cell in 16 out of 200 communities. *T. elliotti* was never detected in any community, a single cell being identified, probably spuriously, to that species. Overall, the four other species (*T. thermophila*, *T. americanis*, *T. borealis*, *T. pyriformis*) were all present in 90% of the samples, and at least three species were detected in the 10% remaining.
